# Genetic underpinnings of sociability in the UK Biobank

**DOI:** 10.1101/781195

**Authors:** J Bralten, CJHM Klemann, NR Mota, W De Witte, C Arango, C Fabbri, MJ Kas, N van der Wee, BWJH Penninx, A Serretti, B Franke, G Poelmans

## Abstract

Difficulties with sociability include a tendency to avoid social contacts and activities, and to prefer being alone rather than being with others. While sociability is a continuously distributed trait in the population, decreased sociability represent a common early manifestation of multiple neuropsychiatric disorders such as Schizophrenia (SCZ), Bipolar Disorder (BP), Major Depressive Disorder (MDD), Autism Spectrum Disorders (ASDs), and Alzheimer’s disease (AD). We aimed to investigate the genetic underpinnings of sociability as a continuous trait in the general population. In this respect, we performed a genome-wide association study (GWAS) using a sociability score based on 4 social functioning-related self-report questions in the UK Biobank sample (n=342,461) to test the effect of individual genetic variants. This was followed by LD score analyses to investigate the genetic correlation with psychiatric disorders (SCZ, BP, MDD, ASDs) and a neurological disorder (AD) as well as related phenotypes (Loneliness and Social Anxiety). The phenotypic data indeed showed that the sociability score was decreased in individuals with ASD, (probable) MDD, BP and SCZ, but not in individuals with AD. Our GWAS showed 604 genome-wide significant SNPs, coming from 18 independent loci (SNP-based h2=0.06). Genetic correlation analyses showed significant correlations with SCZ (rg=0.15, p=9.8e-23), MDD (rg=0.68, p=6.6e-248) and ASDs (rg=0.27, p=4.5e-28), but no correlation with BP (rg=0.01, p=0.45) or AD (rg=0.04, p=0.55). Our sociability trait was also genetically correlated with Loneliness (rg=0.45, p=2.4e-8) and Social Anxiety (rg=0.48, p=0.002). Our study shows that there is a significant genetic component to variation in population levels of sociability, which is relevant to some psychiatric disorders (SCZ, MDD, ASDs), but not to BP and AD.

## INTRODUCTION

Sociability, the quality or state of inclining to seek or enjoy companionship, is a human trait which shows significant variability and is continuously distributed in the general population (Reeb-Sutherland, Levitt, & Fox, 2012). Difficulties with this trait include a tendency to avoid social contacts and activities, and to prefer being alone rather than being with others, potentially linked with a desire to avoid social embarrassment and has been associated with risks of adverse physical and mental health outcomes (Cacioppo, Grippo, London, Goossens, & Cacioppo, 2015; Holt-Lunstad, Smith, Baker, Harris, & Stephenson, 2015) including psychiatric disorders and neurological disorders (Addington, Penn, Woods, Addington, & Perkins, 2008; Bora & Berk, 2016; Dickerson, 2015; Wilson et al., 2007). Avoiding social contacts is an aspect frequently observed in autism spectrum disorders (ASDs), where persistent deficits in social communication and social interaction is one of the core deficits (Lord, Elsabbagh, Baird, & Veenstra-Vanderweele, 2018). Additionally, social withdrawal is a common feature in Major Depressive Disorder (MDD)(Bora & Berk, 2016; Kupferberg, Bicks, & Hasler, 2016), SChiZophrenia (SCZ)(Addington & Addington, 2008; Galderisi, Mucci, Buchanan, & Arango, 2018; Green, Horan, & Lee, 2015) and BiPolar disorder (BP)(Filizer, Cerit, Tuzun, & Aker, 2016) and also observed in neurological disorders like Alzheimer’s disease (AD)(Dickerson, 2015; Reichman & Negron, 2001; Winograd-Gurvich, Fitzgerald, Georgiou-Karistianis, Bradshaw, & White, 2006). Difficulties in sociability can therefore potentially be seen as a cross-disorder overlapping trait in these frequently co-occurring disorders (i.e. (Chisholm, Lin, Abu-Akel, & Wood, 2015; Lee & Lyketsos, 2003; Rai et al., 2018)) with potentially shared underlying mechanisms relating them. The withdrawal from friends, family, and colleagues, is one of the symptoms that is particularly burdensome for relatives of those affected by such disorders. Interestingly this withdrawal from social events and social relationships is one of the earliest signs of disease in groups of patients, suggesting it to be a potential target phenotype for early preventive intervention (Cross, Scott, & Hickie, 2017; Nelis et al., 2011).

Psychiatric and neurological disorders that share difficulties with sociability as a common feature are known to be heritable (estimates are ~64-91% for ASD (Sandin et al., 2017), ~30-40% for MDD (Sullivan, Neale, & Kendler, 2000), ~79% for SCZ (Hilker et al., 2018), ~60% for BP (Johansson, Kuja-Halkola, Cannon, Hultman, & Hedman, 2019), ~60-80 for AD (Gatz et al., 2006). Genome-wide association studies (GWAS) are the current gold standard for localizing common genetic risk variants important for these disorders, comparing people with the disorder (cases) to people without the disorder (controls). Results have been mixed, and extremely large sample sizes were needed to find robust genetic risk variants (ASD: n= 18381 cases and 27969 controls, 5 loci (Grove et al., 2019), MDD: n=246363 cases and 561190 controls, 102 loci (D. Howard et al., 2019), SCZ: n=36,989 cases and 113,075 controls, 108 loci (Ripke et al., 2014), BP: n= 20,352 cases and 31,358 controls, 30 loci (Stahl et al., 2019), AD: n=71,880 AD and AD-by-proxy and 383378 controls, 29 loci (Jansen et al., 2019)). A limitation in the case control design of these studies is the well-known phenotypic heterogeneity observed in these disorders, therefore ‘cases’ can consist of individuals with very diverse sets of symptoms, severity and clinical course. Next to that, comorbidity between disorders and overlap in symptomatology between disorders makes it difficult to select a group of ‘cases’ that includes just one disorder. This makes it, despite the high prevalence of these disorders, difficult to select cases for which genetic material for GWAS is available. Additionally, recent genetic evidence from the Brainstorm Consortium shows that psychiatric disorders share common genetic risk factors (Anttila et al., 2018) and diagnostic criteria do not follow the underlying biology of these disorders well. Investigating the genetics of common overlapping symptoms between disorders might therefore be a good approach to understand the underlying biological mechanisms involved in these disorders, and the co-morbidity of disorders.

Sociability is a multi-determined complex behaviour, which can be modulated by several factors – such as, among others, temperament, personality traits, disability status, aging, social environment and socio-economic status (Holt-Lunstad et al., 2015). There is also growing evidence that, at least in part, sociability is an independent behavioural trait with a specific biological substrate at its basis (Bellack et al., 2004; Kitamura & Suga, 1991; Reichman & Negron, 2001). Indeed the aspects that compose sociability, like loneliness and social interaction have been shown to be moderately to highly heritable by twin and family studies (Boomsma, Cacioppo, Slagboom, & Posthuma, 2006; Boomsma, Willemsen, Dolan, Hawkley, & Cacioppo, 2005; Distel et al., 2010; Ordonana et al., 2013; Stein et al., 2017). Genome-wide association studies investigating the influence of common genetic risk factors involved in loneliness (Gao et al., 2017) and social interaction and isolation (Day, Ong, & Perry, 2018) indeed showed their involvement as well as genetic links to depressive symptoms. However, to our knowledge the underlying biological basis of sociability, as a composed construct including loneliness, social interaction, social isolation and social embarrassment is still to be identified and may differ between diseases, but also between diseases and healthy subjects that manifest this trait.

With difficulties in sociability being a common feature of multiple disorders (Kas et al., 2019; Porcelli et al., 2019), being distributed throughout the population and having a biological basis, studying the genetic underpinnings of sociability in the general population can be an elegant alternative way to study the genetics of overlapping symptoms. The recent increase in genotyped population cohorts, in particular the UK Biobank (Bycroft et al., 2018; Sudlow et al., 2015), increased GWAS sample sizes to hundreds of thousands of individuals. These sample sizes provide the statistical power to detect genetic variants with small effect sizes that are likely to contribute to complex phenotypes, like sociability. In the current study we aim to investigate the genetic contribution to sociability in the general population and if there is a genetic link to the psychiatric disorders ASD, MDD, BP and SCZ as well as the neurological disorder AD. In this light we studied an aggregated construct composed of four sociability-related self-reported questions in the UK Biobank sample and investigated single variant genetic association as well as phenotypic and genetic overlap with the aforementioned disorders.

## METHODS

### Subjects

The UK Biobank (UKBB) is a major population-based study, which includes more than 500,000 individuals aged between 37 and 73 years. The initial data is collected between 2006 and 2010 and includes questionnaire data, genetic data and neuroimaging data. The study design and sample characteristics of the UK Biobank (UKBB) (http://www.ukbiobank.ac.uk/about-biobankuk/) have been extensively described elsewhere (Bycroft et al., 2018; Sudlow et al., 2015). The UKBB was approved by the National Research Ethics Service Committee North West Multi-Centre Haydock and all participants provided written informed consent to participate in the study.

### Sociability phenotype

Four questions in the UK Biobank questionnaire were selected based on their link to sociability (1. Frequency of friend / family visits, 2. Leisure /social activities, 3. Worry after social embarrassment, 4. Loneliness). Based on the answers to these four questions we constructed a sociability measure, where the total score per participant is a sum of the scores on the questions. The sociability score was scored in such a way that a higher score indicated more difficulties with sociability (see **Supplement 1**). Participants were excluded if they had gross somatic problems which could be related to social withdrawal (BMI < 15 or BMI > 40, narcolepsy (all the time), stroke, tinnitus, deafness or brain related cancers) or if they answered that they had “No friends/family outside household” to the question in data-field ‘frequency of family/friends visit’ or “Do not know” or “Prefer not to answer” to any of the questions. We grouped individuals with psychiatric and neurological disorders of interest. For MDD individuals were grouped by ‘probable MDD’ and the occurrence of ‘Recurrent Depressive Episodes’ (D. M. Howard et al., 2018), for SCZ individuals were grouped that had a Schizophrenia diagnosis, for AD individuals were grouped with an Alzheimer diagnosis, for Autism Spectrum Disorders (ASD) individuals were grouped that had childhood autism, atypical autism or Asperger, and for Bipolar Disorder (BP) all individuals that had a Bipolar Disorder diagnosis were grouped. Mean values and distributions were calculated per group using SPSS and compared between individuals with and without neuropsychiatric disorders using general linear models (correcting for age, sex, and assessment center).

### SNP genotyping and quality control

Genotyping details for UKBB participants have been reported previously (Bycroft et al., 2018). Briefly, 49,950 participants were genotyped using the UK BiLEVE Axiom Array and 438,427 participants were genotyped using UK Biobank Axiom Array. The Haplotype Reference Consortium (HRC) and UK10K was the imputation reference sample. For ethnicity we selected self-reported “white” ancestry and subsequently performed a PCA analyses projecting the UKBB participants onto the 1000 Genome Project principal component coordination and excluded individuals that fell out of the cluster (based on PC1 and PC2). Genetic relatedness was calculated with KING kinship (https://kenhanscombe.github.io/ukbtools/articles/explore-ukb-data.html and http://www.ukbiobank.ac.uk/wp-content/uploads/2014/04/UKBiobank_genotyping_QC_documentation-web.pdf) to identify first and second-degree relatives and subsequently ‘families’ were created and only one individual from created ‘families’ was included on the analysis. Single nucleotide polymorphisms (SNPs) with minor allele frequency <0.005, Hardy-Weinberg equilibrium test P value < 1 × 10-6, missing genotype rate >0.05 and imputation quality of INFO < 0.8 were excluded. In the current study 342.461 participants of European ancestry with both genotype and sociability scores available were selected for further analysis.

### Genome-wide association analysis

Genome-wide association analysis with the imputed marker dosages was performed in PLINK using a linear regression model with the sociability measure as the dependent variable and including sex, age, 10 PCA’s, assessment center and genotype batch as covariates. Robustness analyses were included by running five split-half validation analyses splitting our sample five times in two equally sized groups and comparing single variant results as well as excluding individuals with known psychiatric and neurological disorders. Genome-wide association analyses on the four single questions that comprise our sociability score was performed using linear and logistic regression models correcting for sex, age, 10 PCAS, assessment center and genotype batch. To assess the proportion of phenotypic variance explained by common variants, we applied LD score regression (https://github.com/bulik/ldsc) that estimates heritability attributable to genome-wide SNPs (SNP-based heritability) from our sociability GWAS summary statistics based on the slope of the LD score regression (B. Bulik-Sullivan et al., 2015; B. K. Bulik-Sullivan et al., 2015).

### Genetic correlation analyses

To evaluate the extent of shared common variant genetic architectures between sociability and SCZ, BP, MDD, ASD and AD (Grove et al., 2019; Lambert et al., 2013; Ripke et al., 2014; Stahl et al., 2019; Wray et al., 2018), as well as the traits loneliness and social anxiety (Gao et al., 2017; Stein et al., 2017), the bivariate genetic correlations attributable to genome-wide SNPs (rg) were calculated using LD score regression. The GWAS summary statistics underwent additional filtering steps. Only markers overlapping with HapMap Project Phase 3 SNPs and passing the INFO score ≥0.9 and MAF ≥0.01 filters were included (where available). SNPs with missing values, duplicate rs-numbers, too low a sample size (where available SNPs with an effective sample size less than 0.67 times the 90^th^ percentile of sample size were removed), or that were strand-ambiguous – as well as indels – were removed. We used precalculated LD scores (‘eur_w_ld_chr/’ files; (Finucane et al., 2015)) for each SNP using individuals of European ancestry from the 1000 Genomes project that are suitable for LD score analysis in European populations.

## RESULTS

### Sociability measure

A total of 342.461 adults participants from the UK Biobank complied with the inclusion criteria for our study. The sociability score of all participants is distributed from 0-4, with a mean of 1.3 (see **Supplement 2**). Participants with a psychiatric disorder have a higher mean sociability score than the not affected group (ASD p<0.001, BP<0.001, Probable MDD p<0.001, Recurrent Depressive Episodes p<0.001, SCZ p<0.001), but there was no significant difference between the unaffected group and people with an AD diagnosis (p=0.35)(**Figure 1**).

**Figure 1:**
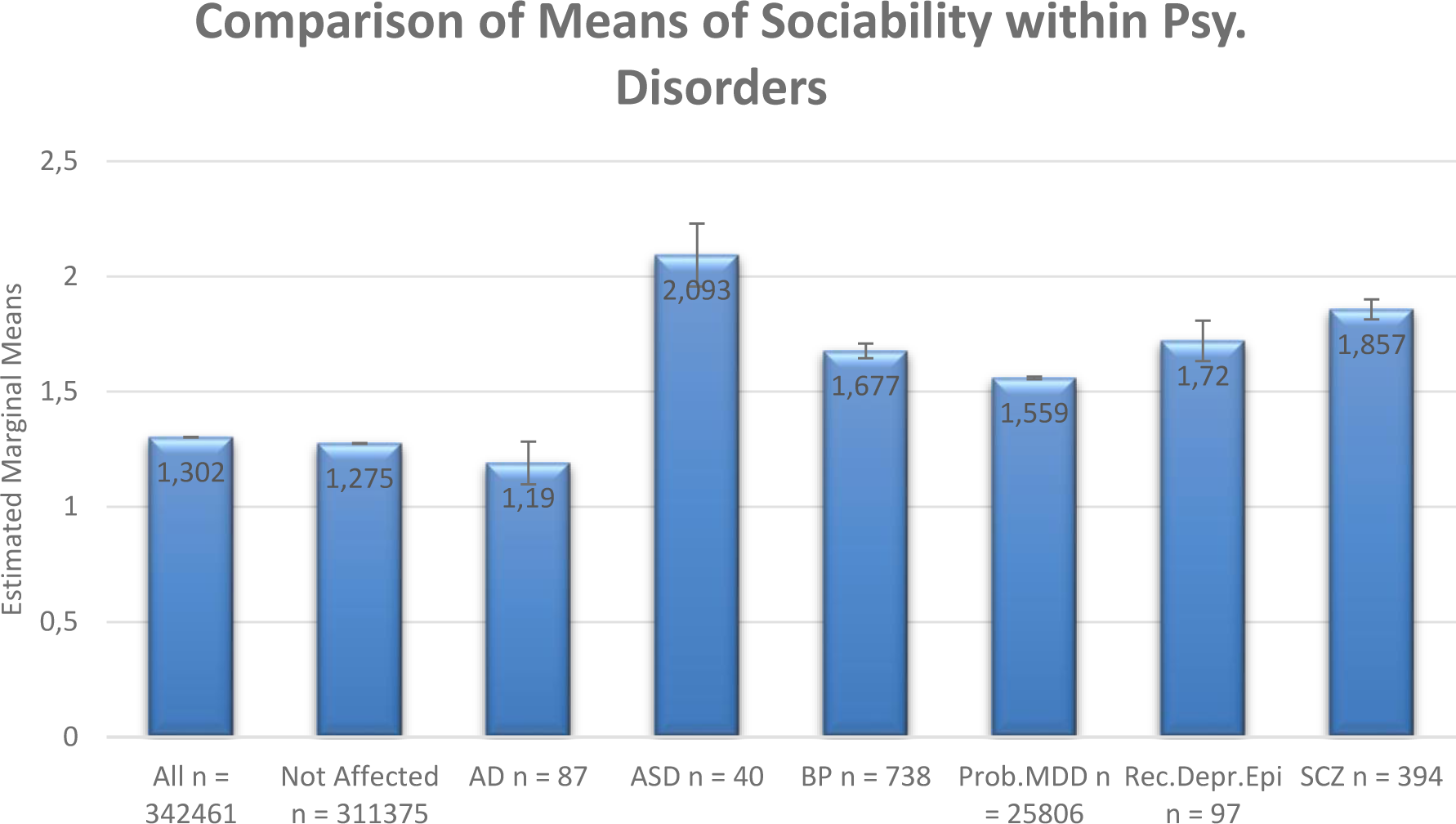
Mean sociability scores in the complete sample (‘All’), individuals without psychiatric and neurological disorders (‘Not Affected’) and with specific disorders of interest (For Alzheimer’s disease (AD) individuals were grouped with an Alzheimer’ diagnosis, for Autism Spectrum Disorders (ASD) individuals were grouped that had childhood autism, atypical autism or Asperger, for Bipolar Disorder (BP) all individuals that had a Bipolar Disorder diagnosis were grouped, for Major Depressive Disorder (MDD) individuals were grouped by ‘probable MDD’ and the occurrence of ‘Recurrent Depressive Episodes’ (Rec. Depr. Epi.), for Schizophrenia (SCZ) all individuals that had a Schizophrenia diagnosis were grouped.

### Genome-wide association analysis

**Figure 2** shows the results of our genome-wide association analyses on the sociability score. In total 604 genetic variants in 18 loci, with 19 lead SNPs surpassed the threshold for genome-wide significance (p<5 * 10-8)(**Table 1**). Our split-half validation showed that all variants had robust associations, showing nominally significant association in all five iterations. Excluding individuals with psychiatric and neurological disorders did not change the results. Single question GWASs showed that all single variant associations had nominal significance in at least two questions. We found that our sociability measure had a SNP-based heritability of 0.0574 (se=0.0025).

**Figure 2:**
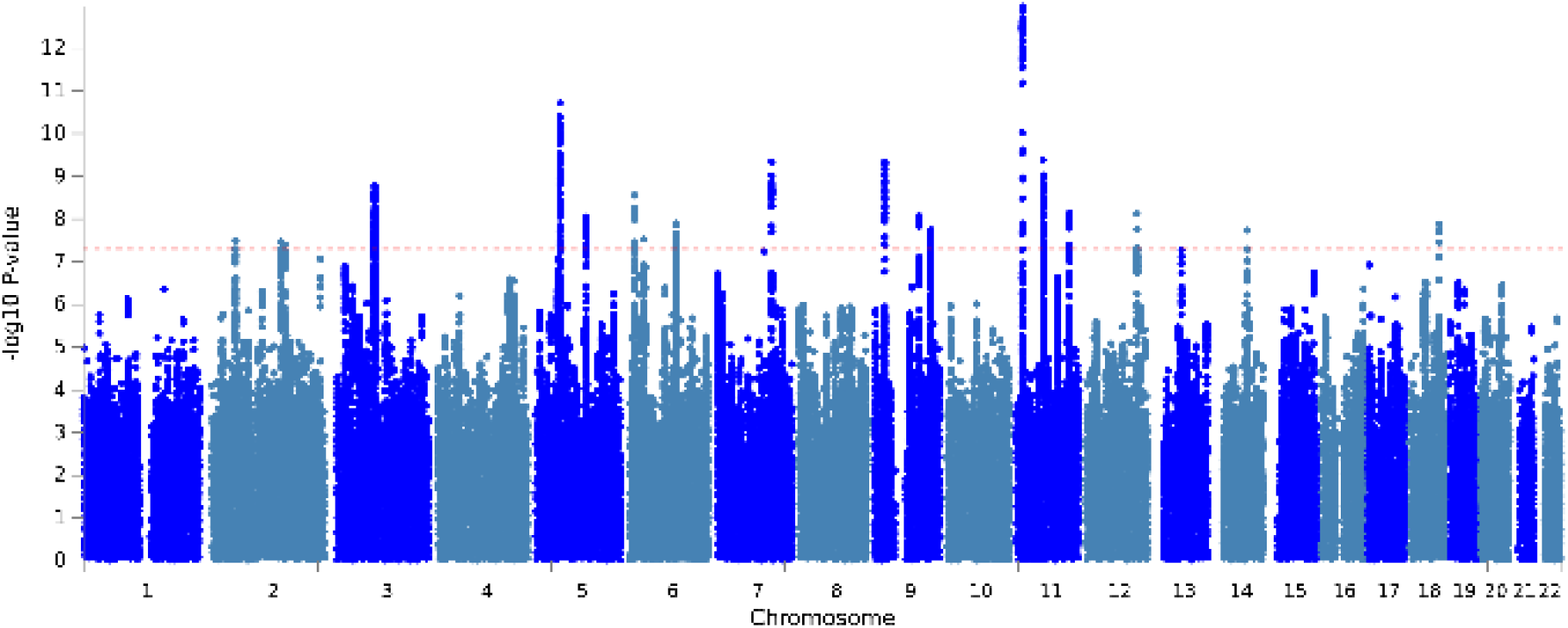
Manhattan plot of the genome-wide association analysis on the sociability construct in the Uk Biobank sample. Every dot indicates the outcome of the linear regression analysis of one single genetic variant with the sociability measure as the dependent variable and including sex, age, 10 PCA’s, assessment center and genotype batch as covariates. On the x-axis you can find the chromosomes, on the y-axis you can find the −log 10 association p-value. The red dotted line indicates the threshold for genome-wide significance.

**Table 1:** Table 1 shows the location, the significance value and the nearest gene for the 19 lead SNPs of the 18 genomic loci that pass the threshold for genome-wide significance (p<5 * 10-8) in the GWAS of the sociability measure in the UK Biobank sample. chr: chromosome, pos: position.

| leadSNP | chr | pos | p-value | Loci | start | end | nearestGene |
| --- | --- | --- | --- | --- | --- | --- | --- |
| **rs10180695** | 2 | 51292332 | 3,19986E-08 | 1 | 51203466 | 51306745 | *AC007682.1* |
| **rs202220108** | 2 | 148910423 | 3,39779E-08 | 2 | 1,49E+08 | 1,49E+08 | *MBD5* |
| **rs57601252** | 2 | 157100293 | 4,08658E-08 | 3 | 1,57E+08 | 1,57E+08 | *NR4A2* |
| **rs4266214** | 3 | 82003207 | 1,61066E-09 | 4 | 81630790 | 82402319 | *RP11-359D24.1* |
| **rs62365541** | 5 | 50905846 | 1,93203E-11 | 5 | 50582954 | 50974050 | *CTD-2335O3.3* |
| **rs4075651** | 5 | 107878588 | 8,80016E-09 | 6 | 1,08E+08 | 1,08E+08 | *RP11-120B7.1* |
| **rs10456089** | 6 | 11959836 | 2,76627E-09 | 7 | 11959836 | 12056771 | *RP11-456H18.1* |
| **rs4839780** | 6 | 100909398 | 1,27664E-08 | 8 | 1,01E+08 | 1,01E+08 | *SIM1* |
| **rs34979551** | 6 | 100848871 | 1,37312E-08 | 8 | 1,01E+08 | 1,01E+08 | *SIM1* |
| **rs6976111** | 7 | 117495667 | 4,62794E-10 | 9 | 1,17E+08 | 1,18E+08 | *CTTNBP2* |
| **rs3793577** | 9 | 23737627 | 4,68151E-10 | 10 | 23720380 | 23741776 | *ELAVL2* |
| **rs10761244** | 9 | 96387592 | 8,36369E-09 | 11 | 96384524 | 96460883 | *PHF2* |
| **rs71507562** | 9 | 121640304 | 1,79161E-08 | 12 | 1,22E+08 | 1,22E+08 | *TUBB4BP6* |
| **rs34588274** | 11 | 13269946 | 1,05085E-13 | 13 | 13268067 | 13350131 | *ARNTL* |
| **rs527528** | 11 | 57433327 | 4,11682E-10 | 14 | 57404779 | 57756568 | *ZDHHC5* |
| **rs4245154** | 11 | 113388674 | 7,36147E-09 | 15 | 1,13E+08 | 1,13E+08 | *DRD2* |
| **rs3742021** | 12 | 109883117 | 7,61756E-09 | 16 | 1,1E+08 | 1,1E+08 | *MYO1H* |
| **rs3784060** | 14 | 72114138 | 1,83262E-08 | 17 | 71757418 | 72162901 | *SIPA1L1* |
| **rs2216270** | 18 | 63656060 | 1,30878E-08 | 18 | 63650084 | 63668364 | *RP11-389J22.3* |

### Genetic correlation

The genetic correlation between sociability and SCZ (0.1532, se=0.0156, p=9.8 * 10-23), MDD (0.6761, se=0.0201, p=6.5 * 10-248) and ASDs (0.2694, se=0.0245, p=4.5 * 10-28) was significantly positive. No significant genetic correlation was found between sociability and BP or AD. Sociability was positively genetically correlated with Loneliness (0.4525, se=0.081, p=2.4 * 10-8) and Social Anxiety (0.4755, se=0.154, p=0.002), see **Table 2**.

**Table 2:** Genetic correlation between the sociability measure in UKBB (So) and case control datasets of Bipolar Disorder (BIP), Schizophrenia (SCZ), Major Depressive Disorder (MDD), Autism Spectrum Disorders (ASD), Alzheimer’s disease (AD), and population-based cohorts on Loneliness and Social Anxiety(SoAnx). Significant genetic correlations are indicated in bold.

|  | <u>Rg (se)</u> | <u>p-value</u> | <u>sample size</u> | <u>PMID</u> |
| --- | --- | --- | --- | --- |
| So-BIP | 0.0142<br>(0.019) | 0.4549 | 20,352 cases, 31,358 controls | 31043756 |
| **So-SCZ** | **0.1532**<br>**(0.0156)** | **9.81<sup>e</sup>-23** | **34,241 cases, 45,604 controls,**<br>**1235 trios** | **25056061** |
| **So-MDD** | **0.6761**<br>**(0.0201)** | **6.57 e-248** | **59,851 cases, 113,154 controls** | **29700475** |
| **So-ASD** | **0.2694**<br>**(0.0245)** | **4.5 e-28** | **18,381 cases, 27,969 controls** | **30804558** |
| So-AD | 0.0392<br>(0.0277) | 0.5747 | 17,008 cases, 37,154 controls | 24162737 |
| **So-**<br>**Loneliness** | **0.4525**<br>**(0.0811)** | **2.4<sup>e</sup>-8** | **7556** | **27629369** |
| **So-SoAnx** | **0.4755**<br>**(0.154)** | **0.002** | **11,268** | **28224735** |

## DISCUSSION

In the current study we show that sociability, as constructed from 4 sociability-related self-report questions in the UKBB, was increased in individuals with ASD, BP, (probable) MDD and SCZ, but not in individuals with AD. Performing a genome-wide association study on a sociability score, we report 604 genome-wide significant SNPs, coming from 18 independent loci. We were able to show that sociability in the general population is genetically correlated with SCZ, MDD and ASDs, but not with BP and AD.

By constructing an aggregated measure of sociability in a large population-based cohort we were able to show that difficulties with sociability are indeed a common feature in multiple disorders. With the UK Biobank cohort collecting data on ‘probable MDD’ and the occurrence of ‘Recurrent Depressive Episodes’ and the diagnoses of ASD, BP and SCZ we could clearly show that our construct was increased in these individuals, compared to the individuals who did not report such incidents, and could therefore be considered a cross-disorder feature. Although difficulties with sociability have also been linked to the neurological disorder AD, we did not find a phenotypic link between our sociability construct and AD, even though the AD measure in the UKBB has been reported to capture AD risk in previous genetic studies (Jansen et al., 2019; Marioni et al., 2018). Not finding a phenotypic association between sociability and a family history of AD is potentially in line with previous work that highlight that difficulties in sociability might not be an early sign of AD, but rather a risk factor for AD. Where the risk of developing AD was found to be increased in individuals who reported to feel lonely, as well as with lower levels of cognition and more rapid cognitive decline, post-mortem analysis on the individuals that ceased away showed that loneliness scores were unrelated to summary measures of AD pathology, like nerve plaques and tangles, or tissue damaged by lack of blood flow (Wilson et al., 2007). However, a recent study using an in vivo method, measuring an aggregated measure of cortical amyloid burden in cognitive normal elderly individuals was able to show a link between self-reported loneliness and cortical amyloid burden, a putative biomarker of AD, indicating loneliness to be a symptom relevant to preclinical AD (Donovan et al., 2016). Next to loneliness, the social aspect of our construct has been investigated as well in patients with AD, and maintaining or developing a close friendship network has indeed been shown to be beneficial for cognition, however this social aspect was not associated to psychopathology (as measured by 12 neuropsychiatric symptoms, like disinhibition, sleep and depression/dysphoria)(Balouch, Rifaat, Chen, & Tabet, 2019). However, we do need to highlight that with AD being an age-related disorder, the phenotype we used in the current study is noisy, as the individuals contributing to the UKBB might not have reached the age to develop AD.

After relating our construct to multiple disorders phenotypically, we were interested to measure if our construct was influenced by common genetic variants. Indeed we found a SNP-based heritability of 6%, which is in line with SNP-based heritabilities that have been reported before for related constructs (4.2%-16% for loneliness(Day et al., 2018; Gao et al., 2017), 3.4%-5% for social interactions (Day et al., 2018), 12% for social anxiety (Stein et al., 2017)). By performing a GWAS in the large scale UK Biobank project, we had the statistical power to detect 18 robust genome-wide significant loci, coming from 19 lead SNPs. The strongest association signal was observed on chromosome 11p15 (*rs34588274*, p=1.05E-13). This loci includes the *ARNTL* gene, which is a circadian clock gene (Rudic et al., 2004). Circadian disruptions, like delayed sleep, reduced sleep efficiency, difficulties falling asleep, early awakenings and higher levels of day-time sleepiness are common among SCZ, BP, MDD and neurodegenerative diseases (Landgraf, McCarthy, & Welsh, 2014; Tsuno, Besset, & Ritchie, 2005; Wulff, Gatti, Wettstein, & Foster, 2010). Indeed genetic variants within core clock genes, including *ARTNL* have been linked to psychiatric disorders and AD (Charrier, Olliac, Roubertoux, & Tordjman, 2017; Chen, Peng, Huang, Hu, & Zhang, 2015; Gonzalez et al., 2015; Kim et al., 2015). Interestingly the lead SNP in this region, *rs34588274*, has previously been linked to neuroticism, well-being spectrum (life satisfaction, positive affect, neuroticism, and depressive symptoms, collectively referred to as the well-being spectrum), positive affect and BMI (GWAS catalog, https://www.ebi.ac.uk/gwas/(Baselmans et al., 2019; Nagel et al., 2018)), indicating that our construct indeed seems to capture aspects related to sociability. This is also supported by the significant positive genetic correlations of our GWAS to loneliness (Gao et al., 2017) and social anxiety (Stein et al., 2017).

Also the genome-wide significant loci on chromosome 11q (rs4245154, p=7.36E-09) includes a well-studied candidate gene for multiple disorders, including SCZ, being *DRD2. DRD2*, dopamine receptor 2, encodes the D2 subtype of the dopamine receptor. This is of particular interest because *DRD2* is one of the few examples where GWAS (of SCZ), after increasing sample sizes to the thousands, was able to find genome-wide significance for a previously ‘expected’ candidate gene (Ripke et al., 2014). Since our top SNP was also significant in the GWAS of MDD, as well as the GWAS of SCZ (D. Howard et al., 2019; Ripke et al., 2014), this hints that our sociability construct is tapping onto genetically shared mechanisms between multiple disorders.

Another interesting genome-wide significant loci is the loci on chromosome 9p21 (rs3793577, p=4.68E-10), which includes the *ELAVL2* gene. The protein encoded by this gene is a neural-specific RNA-binding protein, that has also been reported in the latest MDD GWAS (D. Howard et al., 2019).

This is the first study to investigate the genetics of sociability as an added construct, including loneliness, social relationships, social embarrassment and social activities. Although other studies have investigated the influence of common genetic variants on single parts of this construct, like loneliness (Gao et al., 2017), or using a multi-trait (MTAG) design including loneliness, frequency of social interactions, and ability to confide in someone in the same analysis (Day et al., 2018), we went for an added construct. Our construct clearly showed a phenotypic link to multiple disorders, thereby verifying that we are investigating a clinically relevant construct. We did not perform a multi-trait design because our data did not satisfy the assumptions required for performing MTAG analyses, including the genetic correlations between the single questions being higher than 0.7 and a high value of the mean χ^2^-statistic of our trait (Turley et al., 2018).

By performing genetic correlation analyses between our GWAS of sociability and the latest publicly available disorder GWAS summary statistics we were able to show that sociability genetically correlate with SCZ, ASD, and MDD, but not with BP and AD. The finding that the neurological disorder AD does not genetically link to the same construct as psychiatric disorders is in line with previous findings, showing that genetic sharing was high between psychiatric disorders (including significant genetic correlations between ASD & SCZ, SCZ & MDD, SCZ & BP and MDD & BP), but low between AD and psychiatric disorders (ASD, SCZ, MDD and BP all non-significantly correlated with AD) (Anttila et al., 2018). However, the (genetic) relationship between SCZ and BP is well known, making the (genetic) distinction of BP potentially less expected. Although social impairment characterize BP, even during remitted phases, the link between social relationships and BP seems to be relating to depressive symptoms, and not to manic episodes (Goldstein, Miklowitz, & Mullen, 2006). Indeed patients with BP fluctuate between depressive states and manic states, where in the manic states they have periods where they have increased levels of energy and are very sociable. Opposite to the depressive period, the manic periods might be characterized by an increase in social visits, feeling less socially embarrassed and having the energy to visit public social places, the four aspects covered in our sociability construct. This distinction might be in line with our results showing a very high genetic correlation to depression (clinical MDD), that might be diluted in a clinical sample that includes both depressive and manic episodes.

This study should be viewed in the light of some strengths and limitations. A major strength of the current study is the large sample size, providing us the power to detect prosperous single-variant genetic associations as also shown by our robustness analyses. Another main strength is the combined phenotype used, which is capturing a sociability construct shared between disorders. A limitation of the current study is the modest SNP-based heritability found for our sociability measure, indicating we are only touching on part of the heritability of the complex multifactorial disorders showing difficulties with sociability as an overlapping symptom.

To conclude, our data shows there is a significant genetic component to the variation in population levels of sociability. The genetic contribution to sociability is relevant to the psychiatric disorders ASD, MDD and SCZ, but not to BP and the neurological disorder AD.

## ACKNOWLEDGEMENTS

The PRISM project (www.prism-project.eu) leading to this application has received funding from the Innovative Medicines Initiative 2 Joint Undertaking under grant agreement No 115916. This Joint Undertaking receives support from the European Union’s Horizon 2020 research and innovation programme and EFPIA. This publication reflects only the authors’ views neither IMI JU nor EFPIA nor the European Commission are liable for any use that may be made of the information contained therein. Support by the EU H2020 Program under the Innovative Medicines Initiative 2 Joint Undertaking with grant agreement 777394 (AIMS-2-TRIALS), the Spanish Ministry of Science, Innovation and Universities, Instituto de Salud Carlos III (PI14/00397, PI14/02103, PIE16/00055, PI17/00819, PI17/00481), co-financed by ERDF Funds from the European Commission, “A way of making Europe”, CIBERSAM, Madrid Regional Government (B2017/BMD-3740 AGES-CM-2), EU Structural Funds, EU Seventh Framework Program under grant agreement FP7-HEALTH-2013-2.2.1-2-603196 (Project PSYSCAN), Fundación Familia Alonso, Fundación Alicia Koplowitz.

## SUPPLEMENTARY DATA

**Supplement 1:** Scoring of the questions

Total score for sociability=sum of scores on questions 1-4; range: 0-4. The sociability score was scored in such a way that a higher score indicated more difficulties with sociability.

### QUESTIONS and SCORING

#### 1. Frequency of friend / family visits (Data-Field 1031)

How often do you visit friends or family or have them visit you? (Hint: If this varies, please give an average of how often you visit or have had visits in the last year. Include meeting with friends or family in environments outside of the home such as in the park, at a sports field, at a restaurant or pub.)

| ANSWER | SCORE |
| --- | --- |
| 1. Almost daily | 0 |
| 2. 2-4 times a week | 0.2 |
| 3. About once a week | 0.4 |
| 4. About once a month | 0.6 |
| 5. Once every few months | 0.8 |
| 6. Never or almost never | 1 |
| 7. No friends/family outside household | - exclude |
| 8. Do not know | - exclude |
| 9. Prefer not to answer | - exclude |

#### 2. Leisure / social activities (Data-Field 6160)

Which of the following do you attend once a week or more often? (You can select more than one) (Hint: If this varies, please think about activities in the last year.)

| ANSWER | SCORE |
| --- | --- |
| 1. Sports club or gym |  |
| 2. Pub or social club | At least one of these answers: 0 |
| 3. Religious group |  |
| 4. Adult education class |  |
| 5. Other group activity |  |
| 6. None of the above | 1 |
| 7. Prefer not to answer | - exclude |

#### 3. Worry after social embarrassment (Data-Field 2000)

Do you worry too long after an embarrassing experience?

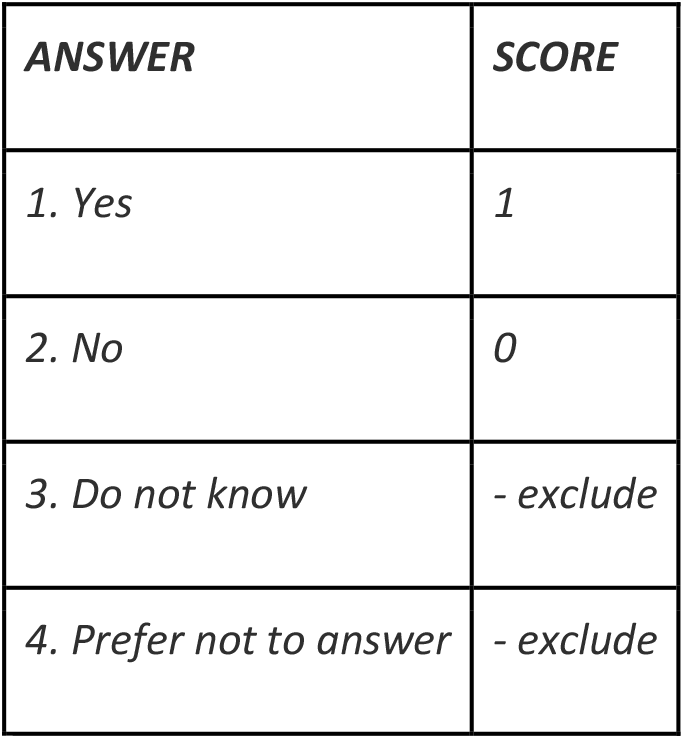

#### 4. Loneliness (Data-Field 2020)

Do you often feel lonely?

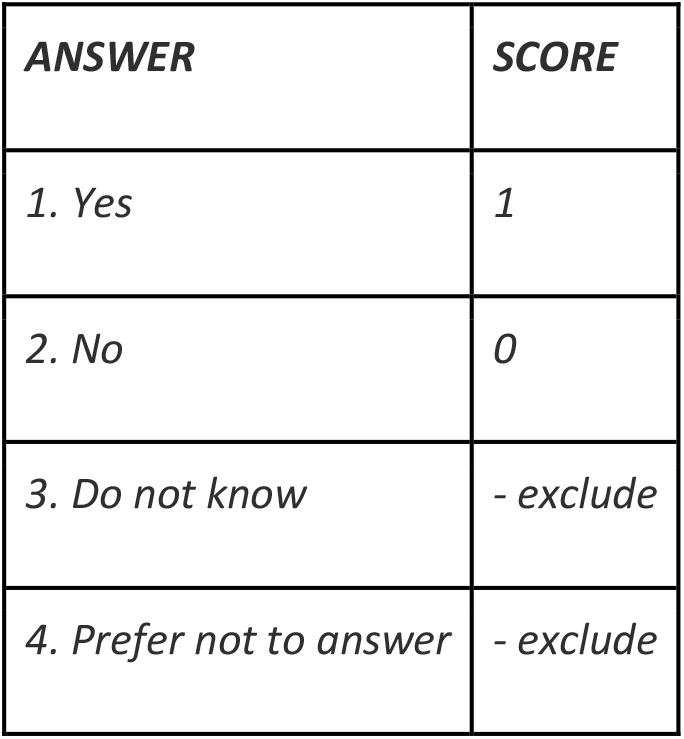

**Supplement 2:** Distribution of total sociability score

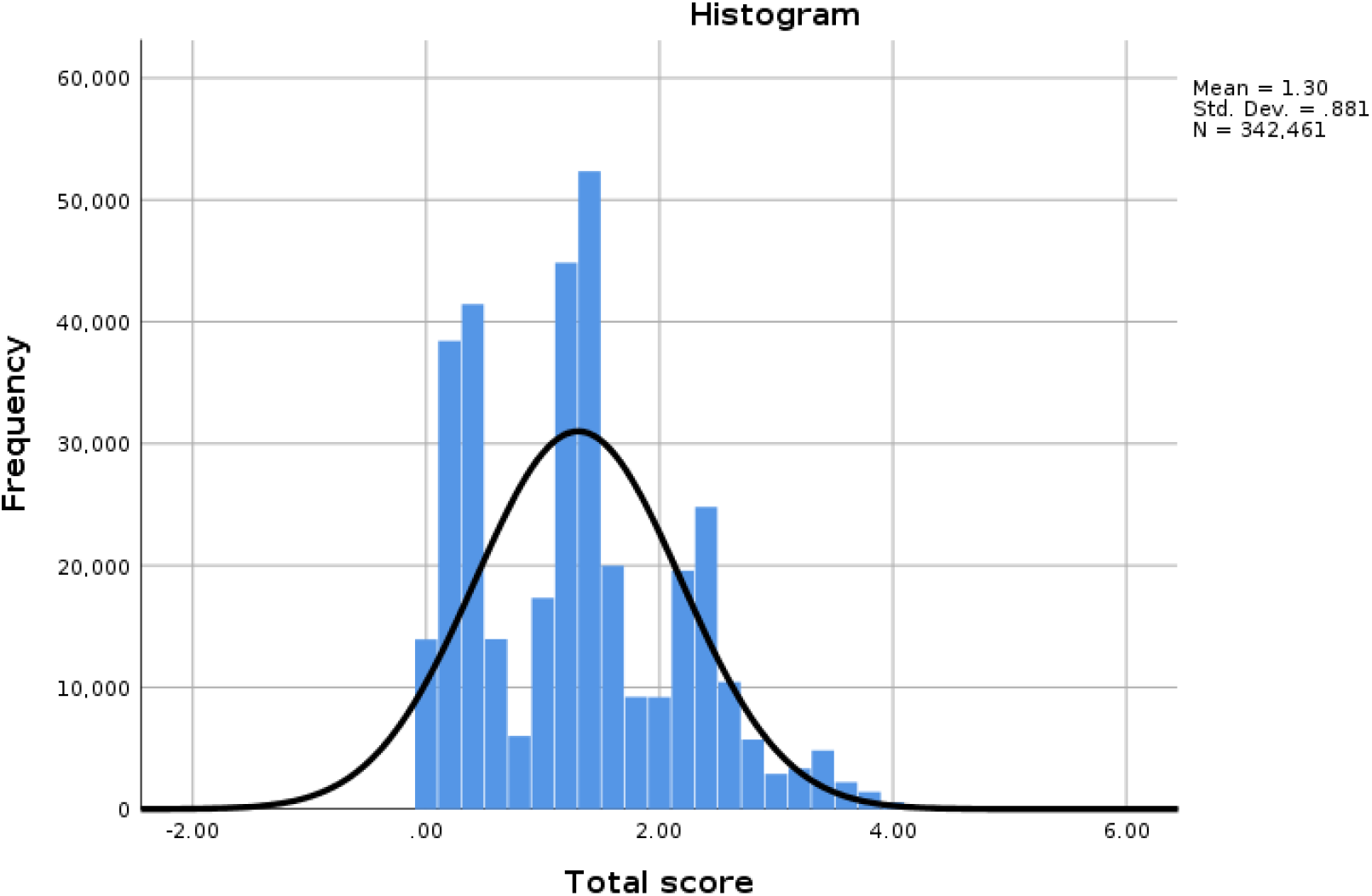

